# Head and eye movement planning differ in access to information during visual search

**DOI:** 10.1101/2022.05.30.493999

**Authors:** Szonya Durant, Tamara Watson

## Abstract

To characterize the process of visual search, reaction time is measured relative to stimulus onset, when the whole search field is presented in view simultaneously. Salient objects are found faster, suggesting that they are detected using peripheral vision (rather than each object being fixated in turn). This work investigated how objects are detected in the periphery when onset in the visual field is due to head movement. Is the process of target detection similarly affected by salience? We test this in 360 degree view with free head and eye movement, using a virtual reality headset with eye tracking. We presented letters and Gabor patches as stimuli in separate experiments. Four clusters were arranged horizontally such that two clusters were visible at onset either side of a fixation cross (near location) while the other two entered the field of view (FoV) when the participant made an appropriate head movement (far location). In both experiments we varied whether the target was less or more salient. We found an interesting discrepancy in that across both tasks and locations the first eye movement to land near a cluster was closer to the salient target, even though salience did not lead to a faster head movement towards a cluster at the far locations. We also found that the planning of head movement changed the landing of gaze position to be targeted more towards the centres of the clusters at the far locations, leading to more accurate initial gaze positions relative to target, regardless of salience. This suggests that the spatial information available for targeting of eye movements within a given FoV is not always available for the planning of head movements and how a target appears in view affects gaze targeting accuracy.

## 1. Introduction

As humans move around in the world, the visual information available to them at any given time is dependent on their head and eye position. They must then rely on this information and any stored representations of their environment to plan their next head and eye position. This work disentangles what information is available and considers that different types of information may be used for planning head movements and eye movements.

When looking around to find something, often what we are searching for may not be within our current field of view (FoV) and we need to move around to find it. These movements include those of the eye, head, and body. If the item is not found within the first ‘look’ we will need to consider how and where to move next. In these circumstances, the appearance of the search target within our FoV occurs because of our motion. Visual search is an extensively researched task, however the experimental procedures almost exclusively involve the search target appearing within the FoV of the participant without their own volition. Exposure is controlled by the experimenter, arguably even if the participant initiates the trial with a key press. The work presented here compares visual search when targets onset within the visual field at the start of an experimenter defined trial, to when they appear within the visual field due to the act of moving the head - to explore the possibility of distinct types of information driving each situation.

Knowledge about how we carry out visual search has mostly been acquired using stimuli that onset in front of a participant who is encouraged not to move their head and to start with their eyes focused on a central location on a computer monitor. This is for good reason as 2D, rectangular, computer displays with a maximum horizontal extent of around 30 degrees of visual angle have been the best technology available to undertake this research. These experiments have allowed us to understand how attention and eye movements operate during visual search within a fixed field of view (FoV), i.e. the possible light array to be sampled stays the same through the trial. Some targets ‘pop out’ relative to surrounding distractor objects and drive a salience-based recognition process that means they are found with very little delay after onset within the visual field with no saccade needed, or within only a single saccade (Binello et al., 1995). These are targets that are differentiated by some basic attribute that makes them extremely salient relative to the distractor objects (e. g. opposing colours). Other objects do not pop out and an effortful search must take place for these to be found. This involves selecting each item/or ‘clumps’ of items (Hulleman & Olivers, 2017) serially in order to inspect them. This search can be guided by the scene we are searching within. For example, a traditional search task involving coloured letters might allow us to exclude all letters except those of a particular colour even though it is the letter identity we are searching for. In that example, it would seem we are able to extract the locations of the appropriate colours without effort and direct attention to those locations in turn to identify the letters. The suggestion is that serial search occurs by establishing a priority map (Wolfe, 2020) of locations to explore based on the attributes of the objects within the scene that can be ascertained within the current functional FoV (i.e. what is clearly visible, a smaller region than the entire FoV). If the task is not a traditional search task, with many singular items randomly arranged within a standard display area, but involves searching within natural scenes, then the priority map may also take into account what we know about the likely location of any particular target object within that scene, i.e. previous knowledge in the form of spatial memory becomes an input, referred to as contextual cueing (Tatler & Land, 2011).

When searching in a display that extends beyond the full extent of the FoV, such that finding targets may require participants to turn their head, we need to consider how previous expectations of locations currently not visible combine with incoming visual information in the priority map, as the FoV changes due to head movement. In particular, we can ask if the different effects of salience described above would still hold in this context - does pop out occur in the same way as soon as the target appears in the FoV?

Visual search with a changing FoV has been less often investigated. To achieve a changeable FoV studies must be carried out in the real world or a simulation of the real world; either through immersive simulators (mixed reality) or virtual reality. Real world driving eye tracking studies can address a type of visual search. Often focussing on stimulus driven and task driven influences on task based visual search, these studies do not explicitly consider how the action of the participant interacts with the likelihood of detecting a target. Although not specifically about visual search, (Doshi & Trivedi, 2012) compared head and gaze shifts in driving, and have suggested head shifts were more associated with top-down task related guidance, part of which might be searching for relevant information in the scene. This may imply the requirement to turn the head during a search task reduces the capacity of a salient stimulus to ‘pop out’ as attentional capacity is devoted to planning a more complex search sequence through unseen space (Crundall, 2005). Similarly, studies of gaze behaviour while playing a range of sports, suggest players position their gaze to make good use of peripheral vision for tracking of important targets, however these studies commonly don’t address how the important target is located in the first instance (Vater et al., 2020).

In terms of virtual reality, (Olk et al., 2018) investigated visual search in a VR environment and found effects transferred from 2D screens in terms of the overall effect on reaction times of target– distractor discriminability. Shioiri et al. (2018) have shown that contextual cueing builds up across trials even when the search display covers 360 degrees and the participant must look around themselves to find the targets. None of these studies carried out in VR have directly addressed whether ability to find a target is changed when it appears within the FoV due to self-motion.

To create an extended visual field, our studies made use of a virtual reality headset to present a 360 degree view of a wrap-around flat (with no stereo or other depth information) image to a seated participant who could rotate their body (and head). In the VR headset there is a restricted FoV as defined by the display of around 110 DVA horizontally and it is this FoV that changes with head movements, as the headset updates in response to head movement.

The task used in this research involves four clusters of stimuli, two of which appear within the FoV at the start of a trial and two of which are beyond the FoV (flanking the initial FoV to the right and to the left, see Fig 1). All four clusters of stimuli are positioned in such a way that the nearest two are in the peripheral visual field when they onset (35 DVA from fixation or the nearest cluster) and are expected to require a head movement so that participants can comfortably respond to the target (they are asked to fixate it). The target stimulus is always present in one of the four clusters, the salience of the target is manipulated. Participants are requested to fixate the target while they register its presence via a key press. Eye movements are recorded while participants carry out the search task to measure reaction time (RT) and the accuracy of the targeting of eye movements. While we did not measure head movements, the position of the stimuli almost guarantees that participants will move their head to fixate the clusters, and any change of FoV is due to head movement. Previous research looking at the coordination of head and eye movements in a VR extended FOV search task showed very few fixation changes beyond 25 degrees are executed without an accompanying head movement (Fang et al., 2015). This is in line with previous research on head and eye coordination (Stern et al., 2005).

**Figure 1:**
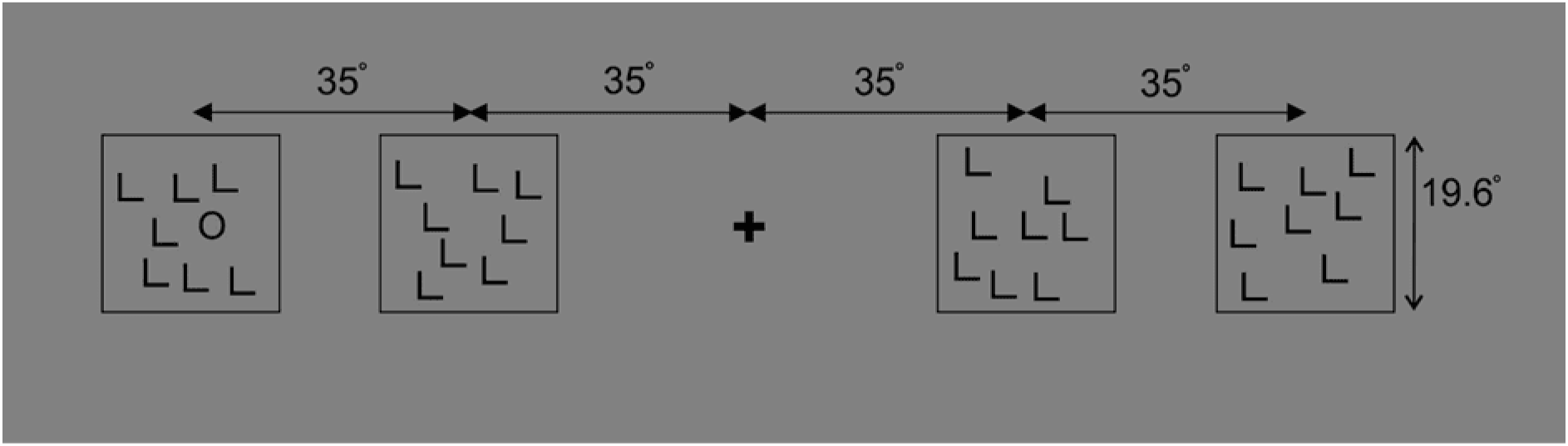
Schematic illustration of the 2D version of the stimulus. 2D images were projected on a sphere using equirectangular projection. Only part of the full 2D image is shown here, the rest was grey. This is an example of the ‘more salient’ condition in which the target is an ‘O’ amongst distractor ‘L’ letters (located here the in far left cluster).

There are, therefore, several interrelated influences that may determine participants’ performance on this task. As we are interested in the potential effect of the salience of targets as they first appear in the periphery, we consider RT as the time it takes to first move the eyes to the cluster containing the target as well as the time to the final fixation leading to a response. Of course this will take longer when the target is in the far cluster, but we might suppose that the more salient target may speed the eyes towards the cluster whether it is present at onset (and hence eyes move to the correct near cluster straight away) or comes into view upon inspection of the near clusters (whether the participant moves in the correct direction first or not, as soon as the salient stimulus in a far cluster enters the FoV it is expected to be identified and therefore move the gaze to the target cluster faster).

Another possible effect of salience may be that it enables the first gaze position in the cluster to land closer to the target. This depends on whether the identity of the stimulus is encoded along with its location. Some research has suggested that under certain conditions in the periphery the compulsory averaging of features occurs, causing positional uncertainty (Greenwood et al., 2009), so speeded reaction times may not necessarily be coupled with increased landing accuracy.

Thus, we consider if any effect of speeding up due to salience observed in reaching the near clusters is reflected in reaction times in reaching the far clusters and similarly if any effect of salience on accuracy of gaze location relative to target is shown at the near and the far clusters in a similar way. Our design disentangles the information available for deciding which cluster to move towards (the direction of the head movement) from that of where to first position the eyes when landing in the cluster (the eye movement to make, given a choice of head movement).

## 2. Methods

We used a HTC Vive (vive.com) Virtual Reality-Head Mounted Display (VR-HMD) capable of collecting gaze data, wired to a PC (Lenovo ThinkStation P510). The software used to present stimuli and record data was Tobii Pro Lab ver. 1.130. The HMD display resolution was 1080 × 1200 pixels per eye, 110 DVA horizontal FoV, refreshing at 90 Hz, with a gaze output frequency of 120 Hz. The Tobii eye tracker was retro-fitted to the HTC Vive headset and eye movements were recorded via the Tobii Pro Lab software. All participants had >95% gaze samples recorded. The raw gaze data was filtered using the default Tobii Pro Lab I-VT fixation filter. The time of a fixation was the first time point at which a fixation began.

### 2.1 Experiment 1 – Letter stimuli

We preregistered our design and methods for Experiment 1 prior to data collection at https://osf.io/umx76/?view_only=062c49054bb046bda94d63c198366ce5.

#### Display and eye tracking

Letter stimuli were drawn in 2D on a 3840 × 1920 pixel image which is presented by Tobii Pro Lab software as a 360 image using equirectangular projection in the HTC Vive headset display (see Display and eye tracking above). This means the letters are drawn on a sphere where 3820 pixels = 360° (degrees of visual angle). The letters were all 3.3° width x 3.6° height, with a stroke thickness of 0.5° (the letters O were anti-aliased to appear smooth). Letters were black on a grey background set at approximately half the max display brightness, but the display was not gamma corrected. The letters were clearly visible when looked at directly.

Four clusters of letters containing 8 letters each were presented centred at −70° (far), −35°(near), 35° (near) and 70° (far) horizontally from the central fixation cross. The two near clusters were visible in peripheral vision while fixating the central fixation cross, while the far clusters required the participant to move their head to bring them into view. Only two clusters could be seen within the FoV. The ‘letter centre’ locations within each cluster were randomly chosen to lie within a square of 19.6° width centred on the cluster location. Each letter was positioned such that the centre of the square containing the letter was at least 5° away from any other letter’s centre. All but one of the letters was an L, the target was either an O (more salient) or a T (less salient). The target was randomly chosen to be one of the locations within the target cluster (replacing one of the L letters). See Figure 1.

Within one experimental run the target appeared in each of the clusters 10 times, yielding 2 (O/T) x 4 (location) x 10 = 80 trials. This resulted in 20 measurements per condition (near/far x O/T). 5 sets of full experimental stimuli were pre-generated and each participant saw one set of these sets. The order of stimuli was randomly shuffled for each participant.

#### Participants

34 participants were recruited through a combination of personal approach, and online advertising to gain course credit/or a £5 reimbursement. 8 were excluded according to data quality criteria (see below), leaving 26. According to (Brysbaert, 2019) for 80% power and a medium effect size (dz = 0.4) for a 2 × 2 repeated measures ANOVA 27 participants are required. Of these 15 were male, 11 female, with a mean age of 20.2 (range 18-25) (one participant did not give their age). All had corrected to normal vision and used glasses or contact lenses in the headset when required.

#### Procedure

After giving consent (the experiment was approved by the RHUL Ethics Committee) and also being given verbal instructions, the participants were seated on a standard office swivel chair and the headset was fitted. They were told they could swivel the chair round.

Participants were given a practise session in which 4 trials were shown, one each O/T, near and far. These practise images were not used in the experiment.

At the start of the experiment, after a 5 point eye tracker calibration, four crosses were shown and the participant was as asked to look at each in turn (in no particular order). This was partially to make sure the participant was facing in the correct direction, so that the fixation cross would appear in front of them and was also used as a calibration check. The central fixation cross remained on screen in between each trial for 3s. Their instructions were: “As soon as a new stimulus appears, look for the ‘O’ or ‘T;. You will not know on each trial whether it is the ‘O’ or ‘T that will be the target. When you have found it, make sure to look at it and then press the button response. Then look back at the fixation cross.” At the end of the experiment the 4 crosses appeared again with the same instruction as at the start. The experimenter remained present and could see the how the stimulus looked to the participant on a monitor screen as well as see their real-time gaze location and the Tobii Pro lab visual indicator of how well the eyes were being tracked.

#### Data checking and analysis

Trials were excluded if the fixation at the start of the trial was further than 9.4° away from the fixation cross. Each fixation was assigned a cluster location according to which of the elements (letters) within the cluster was nearest. If the final three fixations before responding were not in the cluster containing the target, or the participant never fixated within 9.4° of an element, these trials were also excluded. Participants with more than 10% of trials excluded were fully excluded.

Reaction times were calculated in several ways. 1. The time from the appearance of the stimulus to the button press. 2. The onset time of the last fixation in the target cluster (how long it took to decide to respond correctly) to the time a response was recorded. 3. the time of the first fixation in the cluster containing the target (how long it took to decide to look in the correct cluster).

We also calculated the position of the first fixation in each cluster. This was calculated as the first fixation location whose closest element was from that cluster, having moved more than 9.4° from fixation. This was based on observation of the fact that fixations were mostly always on cluster elements.

#### Design

We ran this study as a 2×2 repeated measures, with conditions: location (near/far) and salience (high/low)- analysed for each DV (reaction time in terms of time of first and last fixation in the target cluster; distance from the target of the first fixation in the target cluster) separately.

### 2.2 Experiment 2 – Gabor patch stimuli

We preregistered our design and methods for Experiment 2 after data collection, but prior to analysis at https://osf.io/rf24c/?view_only=3e405aca39f14c7c85fd78b37aa63b98.

#### Stimuli

Gabor stimuli were drawn in 2D on a 1920 × 3840 pixel image which is represented by Tobii Pro Lab software as a 360 image using equirectangular projection in the HTC Vive headset display (see Display and eye tracking above). This means the Gabor patches are drawn on a sphere where 3820 pixels = 360° (degrees of visual angle). The full width at half height of the Gabor patches was 1.7°, their spatial frequency was 0.75 cycles/°. They were drawn on the same grey background as the letters. Their Michelson contrast was calculated as 0.7 based on pixel values, but the display was not gamma corrected. Gabor patches were clearly visible when looked at directly.

Four clusters of Gabor patches were presented centred at −70° (far), −35°(near), 35° (near) and 70° (far) horizontally from the central fixation cross, each containing 8 Gabor patches (see Figure 1). The Gabor patch centre locations within each cluster were randomly chosen to lie within a square of 20.8° width centred on the cluster location. Each Gabor patch’s centre was positioned such as to be at least 6° away from another Gabor patch.

Orientations for the Gabor patches were randomly chosen so that in each cluster half are tilted −70° from target and the other half are 70° from target in the salient condition, and −20°/+20° in the non-salient condition. The target was randomly chosen to be one of the locations within the target cluster, replacing one of the distractor Gabor patches (see Figure 1).

Within one experimental run the target appeared in each of the clusters 5 times, yielding 2 (more salient/less salient) x 2 (vertical/horizontal) x 4 (location) x 5 = 80 trials, with 20 measurements per condition (collapsing vertical and horizontal). 5 sets of full experimental stimuli were pre-generated, each participant saw one set of these, which were randomly shuffled differently for each participant.

#### Participants

26 participants were recruited through a combination of personal approach, and online advertising to gain course credit/or a £5 reimbursement. 3 were excluded due to data quality criteria, leaving 23, see below. According to (Brysbaert, 2019) for 80% power and a medium effect size (dz = 0.4) for a 2 × 2 repeated measures ANOVA, 27 participants are required. Of these 7 were male, the ages were a mean of 25.9 (range 19-45) (one participant did not give their age). All had corrected to normal vision and used glasses or contact lenses in the headset when required.

#### Procedure

After giving consent (the experiment was approved by the RHUL Ethics Committee) and also being given verbal instructions, the participants were seated on a standard office swivel chair and the headset was fitted. They were told they could swivel the chair round. They were given a practise in which four trials were shown, one each horizontal/vertical more salient/less salient, each one at a different cluster location. These practise images were not used in the experiment.

At the start of the experiment, after a 5 point eye tracker calibration, four crosses were shown, which the participant was as asked to look at in turn (in no particular order). This was used as a calibration check. It would also orient the participant to where the central fixation cross would appear. The central fixation cross remained on screen in between each trial for 3s. Their instructions were: “As soon as a new stimulus appears, look for the vertical or horizontal patch. You will not know on each trial whether it is the vertical or horizontal patch that will be the target. When you have found it, make sure to look at it and the press the button response. Then look back at the fixation cross.” The stimulus would disappear when the responded. At the end of the experiment the four crosses appeared again with same instruction as at the start. The experimenter remained present and could see the how the stimulus looked to the participant on a monitor screen as well as see their real-time gaze location and the Tobii Pro lab visual indicator of how well the eyes were being tracked.

#### Data checking and analysis

Trials were excluded if the fixation at the start of the trial was further than 9.4° away from the fixation cross. Each fixation was given a cluster location according to which of the elements was nearest. If none of the fixation cluster locations within the last three before responding were in the cluster containing the target or they never fixated within 9.4° to an element, these trials were also excluded. Participants with more than 10% excluded trials were fully excluded.

We could use the software to log the time from the appearance of the stimulus to the button press. Reaction times were calculated based on eye movements in 2 main ways. We took the time point of the last fixation in the target cluster (how long it took to decide to respond correctly) before responding as one measure and we also took the first fixation in the cluster containing the target as the time to first fixation (how long it took to decide to look in the correct cluster).

We also calculated the position of the first fixation in a cluster. This was calculated as the first fixation location whose closest element was from that cluster, having moved more than 9.4° from fixation. This was based on observation of the fact that fixations were mostly always on cluster elements.

#### Design

We ran this study as a 2×2 repeated measures, with conditions: location (near/far) and salience (high/low)- analysed for each DV (time to first fixation in cluster, time to last fixation in cluster, distance of first fixation in the target cluster form the target) separately.

## 3. Results

### 2.1 Experiment 1 - Letter search task

#### Reaction times

The reaction times we report here are 1. the time it took to first fixate within the target cluster after the onset of the four stimulus clusters. 2. the time of the last fixation within a target cluster before the response was registered relative to the stimulus onset. These were used to assess whether salience enabled the target cluster to be fixated more quickly and whether the effect of salience was the same according to how a cluster appeared in view.

We note that the pattern remains the same between the first and last fixation in the target cluster, with the last fixation in the cluster being around 500ms after the first fixation in the cluster, and salience apparently having the most effect at the near locations. The 500ms delay that is similar across all conditions is possibly due to participants making several fixations on target before pressing the response button. This delay is less on average for the more salient stimuli; 200ms less at the near locations and 75ms at the far locations. This may be due to the initial landing position within the target cluster being closer for more salient targets, which we test below.

A 2 × 2 repeated measures ANOVA was conducted to investigate the effect of near/far location and the salience of the search target on time to first fixation in the target cluster. There was a main effect of location: F_1,25_ = 230.4, p<0.0005 and of target salience F_1,25_ = 67.9, p<0.0005 as well as an interaction F_1,25_ = 51.0, p<0.0005. As expected, letters in the far location took longer to find and the more salient O was quicker to be found than the less salient T, but the interaction indicates (as can be seen when looking at Figure 3a) that the advantage of salience is greatly reduced at the far locations. This means that salient stimuli visible at stimulus onset lead the eyes to move to their cluster faster, but this is not the case for salient stimuli visible only after the participant orients toward one of the near clusters. The pattern of results was the same for the last fixation in the target cluster ahead of response. (Main effect of location: F_1,25_ = 248.4, p<0.0005; main effect of salience: F_1,25_ = 108.7, p<0.0005; interaction: F_1,25_ = 63.4, p<0.0005).

As an explanation for the faster RT to near cluster salient targets we also checked whether the first eye movement after stimulus onset was more often directed toward a near cluster target when it was salient. We found that the chance of immediately looking the correct way if the target was in one of the close clusters was 90.3% (s.d. 7.5%) for the O stimulus and 58.3% (s.d. 13.9%), which was a significant difference T_25_ = 12.3, p<0.0005.

This means that when a more salient target appeared in the visual field due to stimulus onset at the near location it was able to bias the eye movements such that gaze location moved there faster. On the other hand, when a salient target appeared in the visual field during the course of the trial (at the far locations) it was no longer able to bias gaze such that it reached the target cluster faster.

#### Location of the first fixation in a target cluster

We also wished to assess whether the salience of the target had an effect on where a first fixation landed in the target cluster – i. e. to what extent the location of the more salient target was represented whilst the cluster was not in central vision, relative to the less salient target. We recorded the location of the first fixation within the cluster containing the target.

In Figure 2 we see that the largest average distance between the first fixation in the cluster and the target is around 8 degrees, which is around half of the cluster diameter. This is what we would expect if the landing location were random or if always landing in the middle of the cluster. This distance is smaller in some conditions.

**Figure 2:**
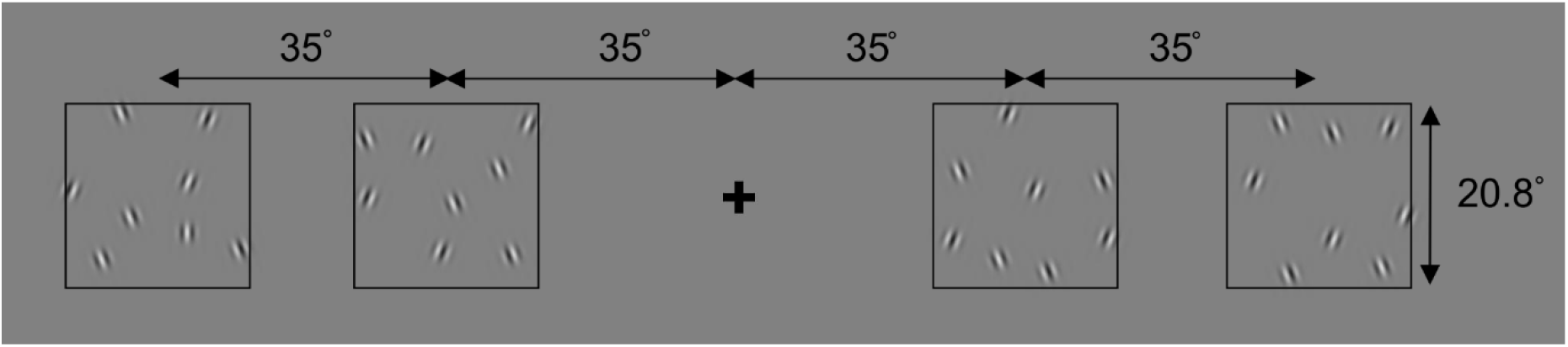
Schematic illustration of the 2D version of the stimulus that was then projected on a sphere using equirectangular projection. Not the full 2D image is shown here, the rest was grey. This is an example of the ‘less salient’ condition, in which the distractors are oriented at +/- 20° from vertical, and the target is vertical (located here in the far left cluster).

A 2 × 2 repeated measures ANOVA was conducted on the effect of near/far location and the salience of the search target on the distance from the target of the first fixation in the cluster containing the target. We found a main effect of location F_1,25_ = 27.0, p<0.0005 and salience F_1,25_ = 38.5, p<0.0005, but no interaction F_1,25_ = 1.5, p=0.24. Thus, the first gaze position within a cluster containing the target was significantly closer to the target if the target was in one of the far clusters, and also if the target was the more salient O target. The advantage of the salient target did not differ across locations. Thus, the RT advantage for the more salient target at the near location was also paired with a better representation of its position. Interestingly the salient stimulus’ position was also well represented at the far location despite it not leading to a faster RT (see Figure 3). It is also interesting to note that the way the cluster entered the field of view also had an effect on how well the first fixation in the cluster was targeted relative to the target stimulus, with the far cluster coming into view during the trial conferring an advantage for both types of targets.

**Figure 3.**
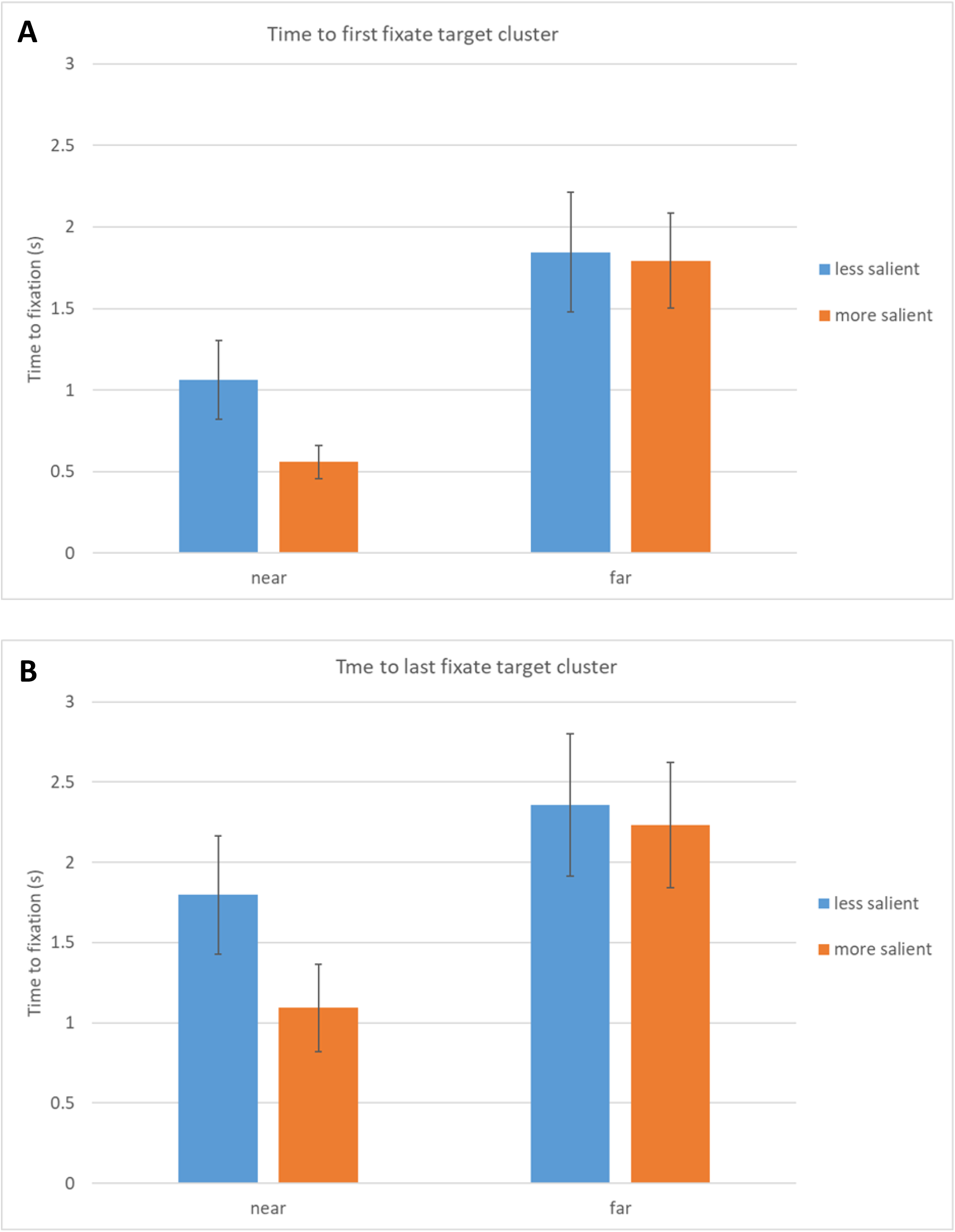
(a) Time to first fixate in the target cluster. (b) time of last fixation in the target cluster. N=26, error bars S.D (note with a repeated measures comparison, error bars do not reflect significance). The less salient target is the ‘T’ and the more salient is the ‘O’ amongst distractor ‘L’ letters. Near and far are the cluster location relative to the fixation cross.

**Figure 4.**
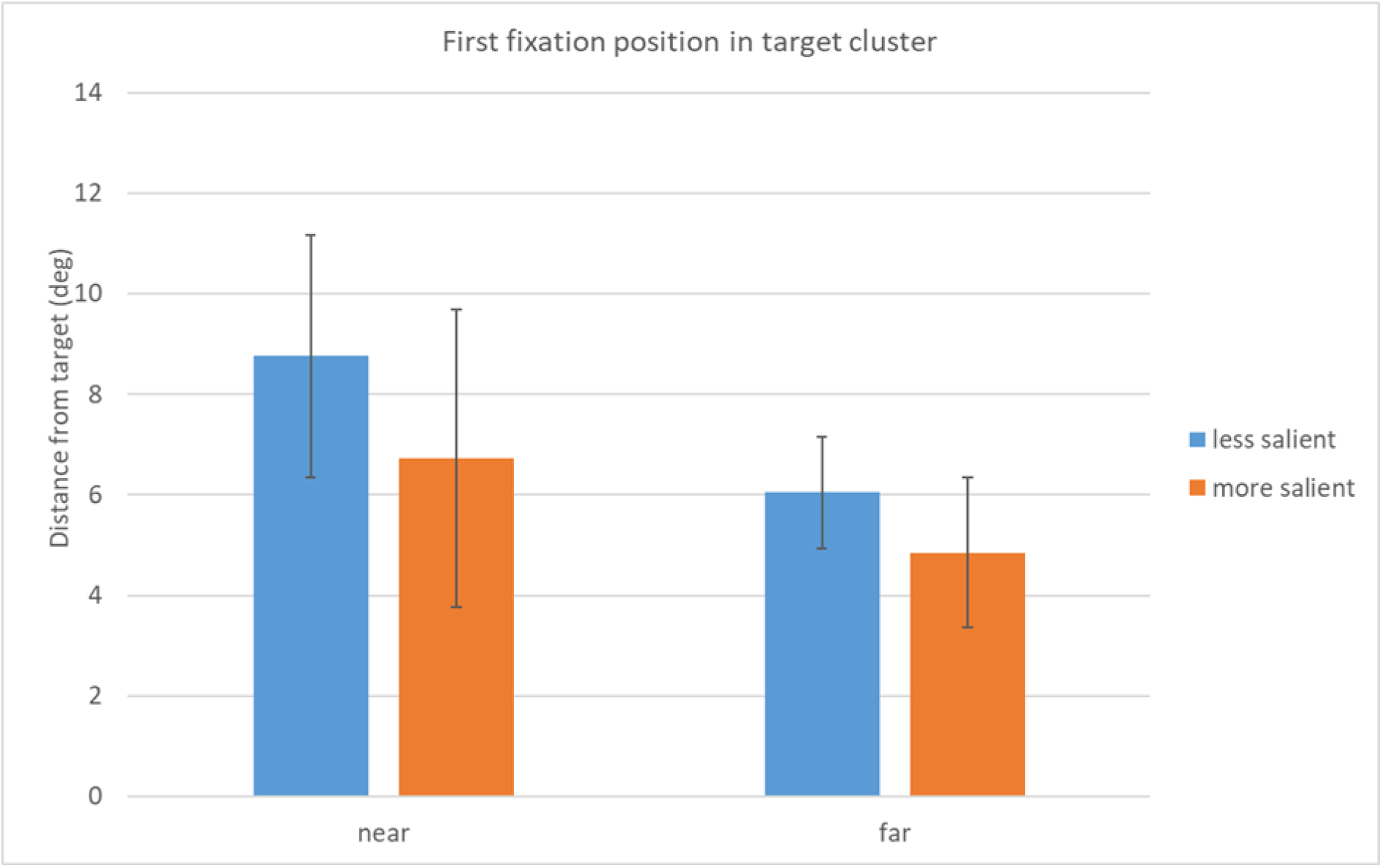
Distance between the first fixation in the target cluster and the target. N=26, error bars s.d. (note with a repeated measures comparison, error bars do not reflect significance). The less salient target is the ‘T’ and the more salient is the ‘O’ amongst distractor ‘L’ letters. Near and far are the cluster location relative to the fixation cross.

‘First fixation’ strategies may be adopted if the participant cannot detect the stimulus from the previous fixation location outside the cluster. One of these would be to target the centre of the cluster, another may be targeting the edge of the cluster closest to the currently fixated cluster. To visualise whether participants show any fixation location bias relative to the location of the target, polar plots were created that show every single first fixation in a target cluster arranged so that polar plot angle is the angle away from the target relative to the centre of the cluster (Fig. 5). Here the centre of the polar plot represents the target, while zero angle represents the direction of the centre of the cluster. The distance is calculated from the first fixation for which the nearest element is in the target cluster. We see that in general the fixations are either on the target or biased towards the centre of the cluster, rather than elsewhere away from the target. This pattern appears to change across conditions where in the far more salient ‘O’ condition; it is clear that the bias is for fixations to be on target, whereas in the far less salient ‘T’ condition it is towards the middle of the cluster. In the near conditions the fixations tend to be more spread out from the centre, suggesting that the strategy here was more often to target the edge of the clusters

**Figure 5.**
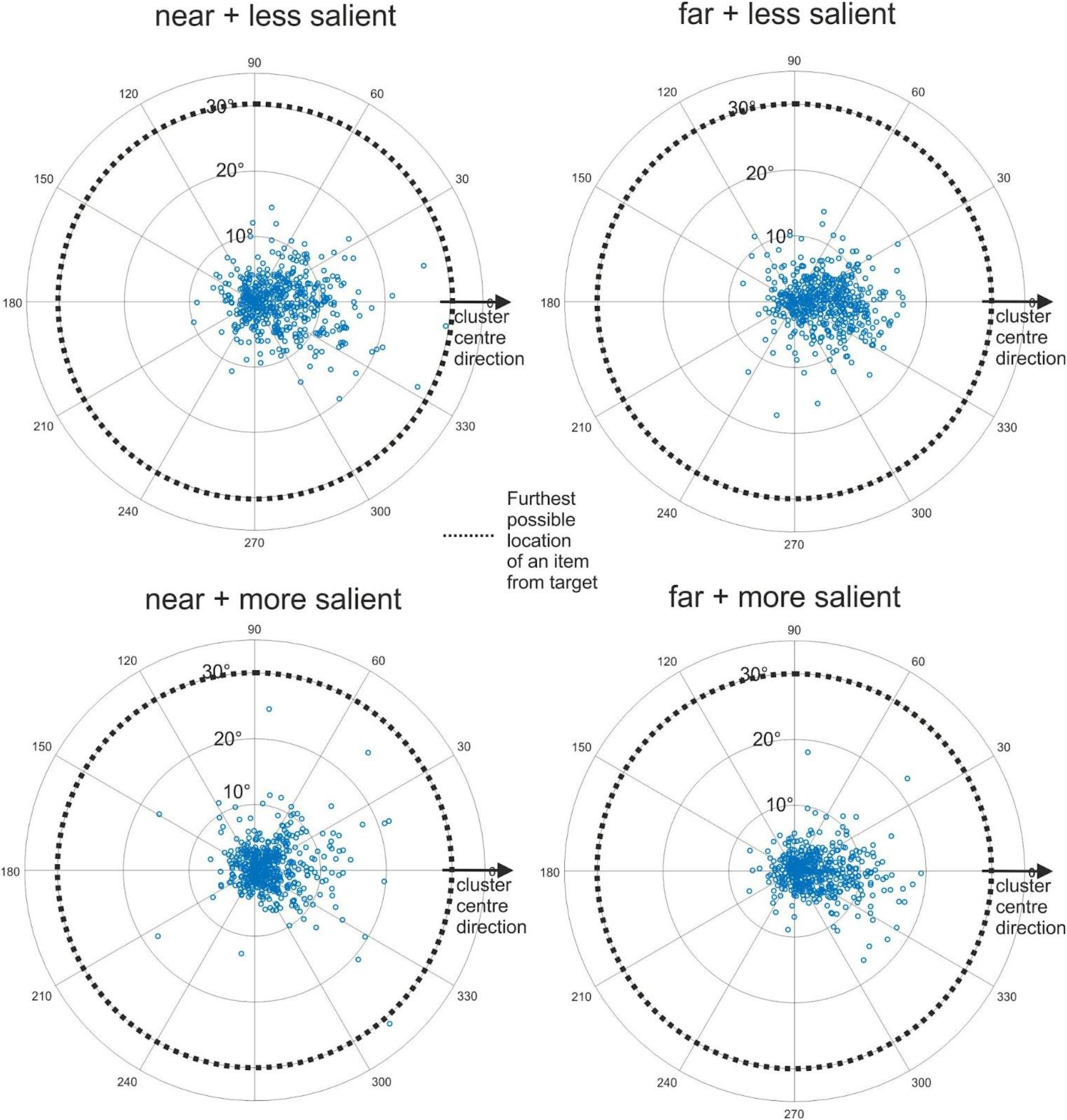
Polar plots of the first fixations in a cluster. The polar plots show every single first fixation in a cluster arranged so that the polar plot angle is the angle away from the target relative to the centre of the cluster.

**Figure 6.**
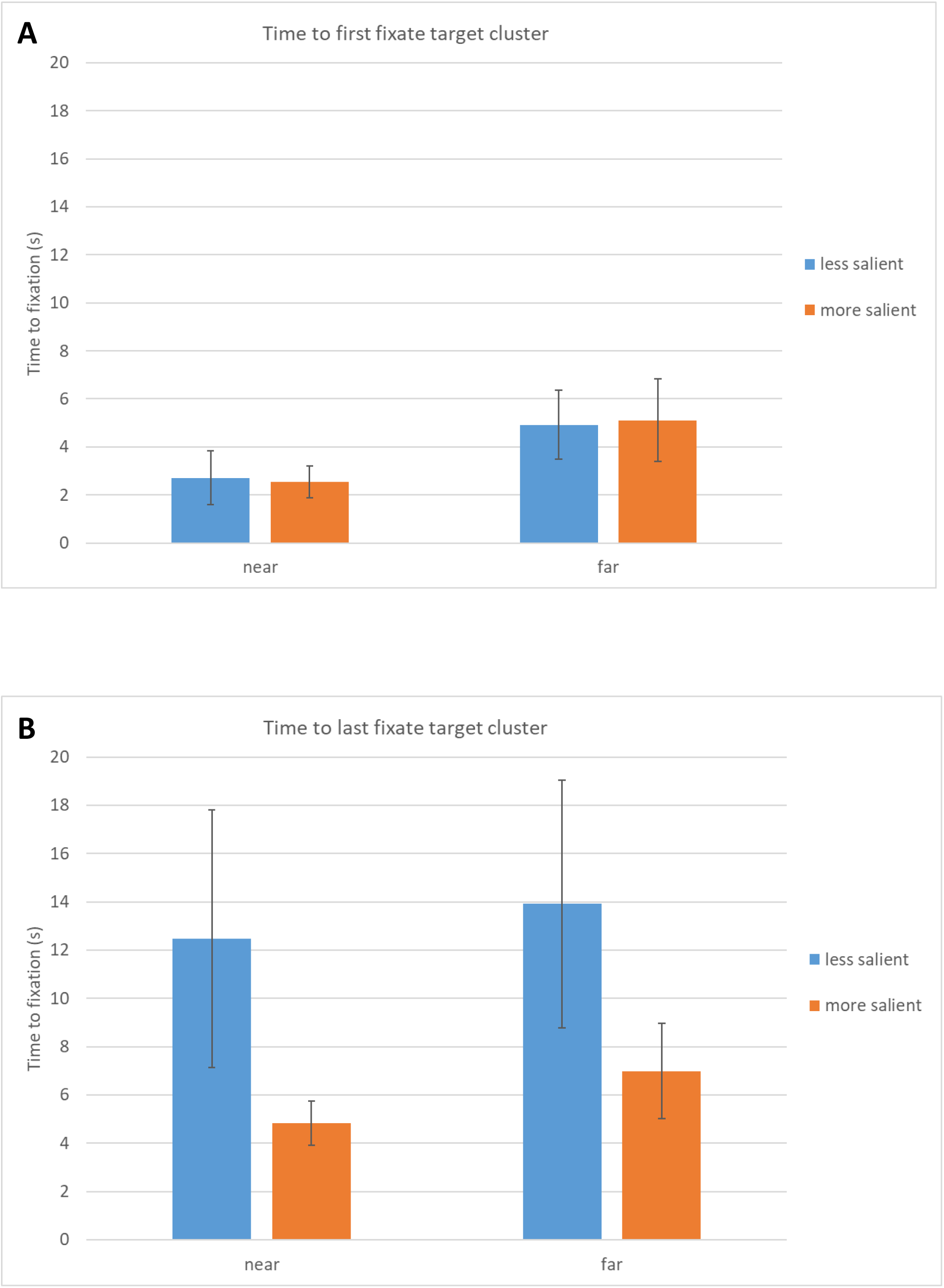
(a) Time to first fixate in the target cluster. (b) time of last fixation in the target cluster. N=23, error bars S.D (note with a repeated measures comparison, error bars do not reflect significance). The less salient target is when distractors are oriented away from the target (horizontal or vertical) stimulus by +/- 20° and in the less salient condition only by +/-110°. Near and far are the cluster location relative to the fixation cross.

If we compare the near to the far plots, we see a similar difference across both the more and less salient conditions; the first fixations in the target clusters are less spread away from the target in the far clusters and more centred around the direction of the centre of the target cluster.

### 3.2 Experiment 2 – Gabor patch search task

The same analysis was applied as was used to investigate responses during the letters task. In this case the more salient stimulus consisted of those stimuli where the distractor Gabor patches were oriented away from the target (horizontal or vertical) stimulus by +/- 110° and in the less salient condition only by +/-20°.

#### Reaction times

As before, the reaction times we report here are 1. the time it took to first fixate within the target cluster after the onset of the four stimulus clusters and 2. the time of the last fixation within a target cluster ahead of response after the stimulus onset. These were used to assess if salience enabled the target cluster to be fixated more quickly and whether the effect of salience was the same according to how a cluster appeared in view.

We note that these two graphs look markedly different from Experiment 1 and there is now a different pattern between the first and last fixation RT in the target cluster. There is a large time difference between first and last fixation. This is around 2s for the more salient stimuli and 9s for the less salient stimuli. This suggests that much more time is needed to decide where the target is after first fixation.

A 2 × 2 repeated measures ANOVA was conducted on the effect of near/far location and the salience of the target on the time to first fixation in the target cluster. There is a main effect of location (F_1,22_ = 93.3, p<0.0001) but no effect of salience (F_1,22_ = 0.01, p=0.92) and no interaction (F_1,22_ = 1.4, p=0.24). As expected, the far target clusters took longer for the gaze to reach, but salience does not speed up the eyes in reaching the cluster. The same analysis was carried out for the last fixation in the target cluster and here we find a main effect of location (F_1,22_ = 14.9, p<0.005) and salience (F_1,22_ = 73.0, p<0.001) and no interaction (F_1,22_ = 0.90, p=0.35. The effect of salience here seems to be driven by the decision time needed after reaching the target cluster for the first time.

This was further confirmed by considering the direction of the first eye movement from fixation. We found that the chance of immediately looking the correct way if the target was in one of the close clusters was 49.8% (s.d. 11.0%) for the more salient stimulus and 47.7% (s.d. 13.4%) for the less salient stimulus, which was not significantly different (T_20_ = 0.45, p=0.65). Unlike in the letters task, in this task the target was not salient enough to influence the direction of the first eye movement. This result accords with the observation that this task was very difficult and even the most salient targets did not ‘pop-out’. They seemed to require a kind of serial consideration of stimuli before a confident response could be made. The longer response times in this task (fastest condition 3 seconds) relative to the letter task (fastest condition 0.6 seconds) also reflect this.

#### Location of the first fixation in a target cluster

As before, the distance from the target of the first fixation near a cluster was assessed (see Figure 7). A 2 × 2 repeated measures ANOVA was conducted on the effect of near/far location and target salience on the distance from the target of the first fixation in the cluster containing the target. As for the letter task, there was a main effect of location (F_1,22_=12.5, p<0.005) and salience (F_1,22_=83.1, p<0.0005), and again no interaction (F_1,22_=0.05, p=0.83). This result suggests that despite the participants being at chance regarding whether they moved their gaze initially in the direction of the more salient target stimulus, the more salient targets still provided some location information to be more reliably targeted - whether they were in the near or far cluster.

**Figure 7.**
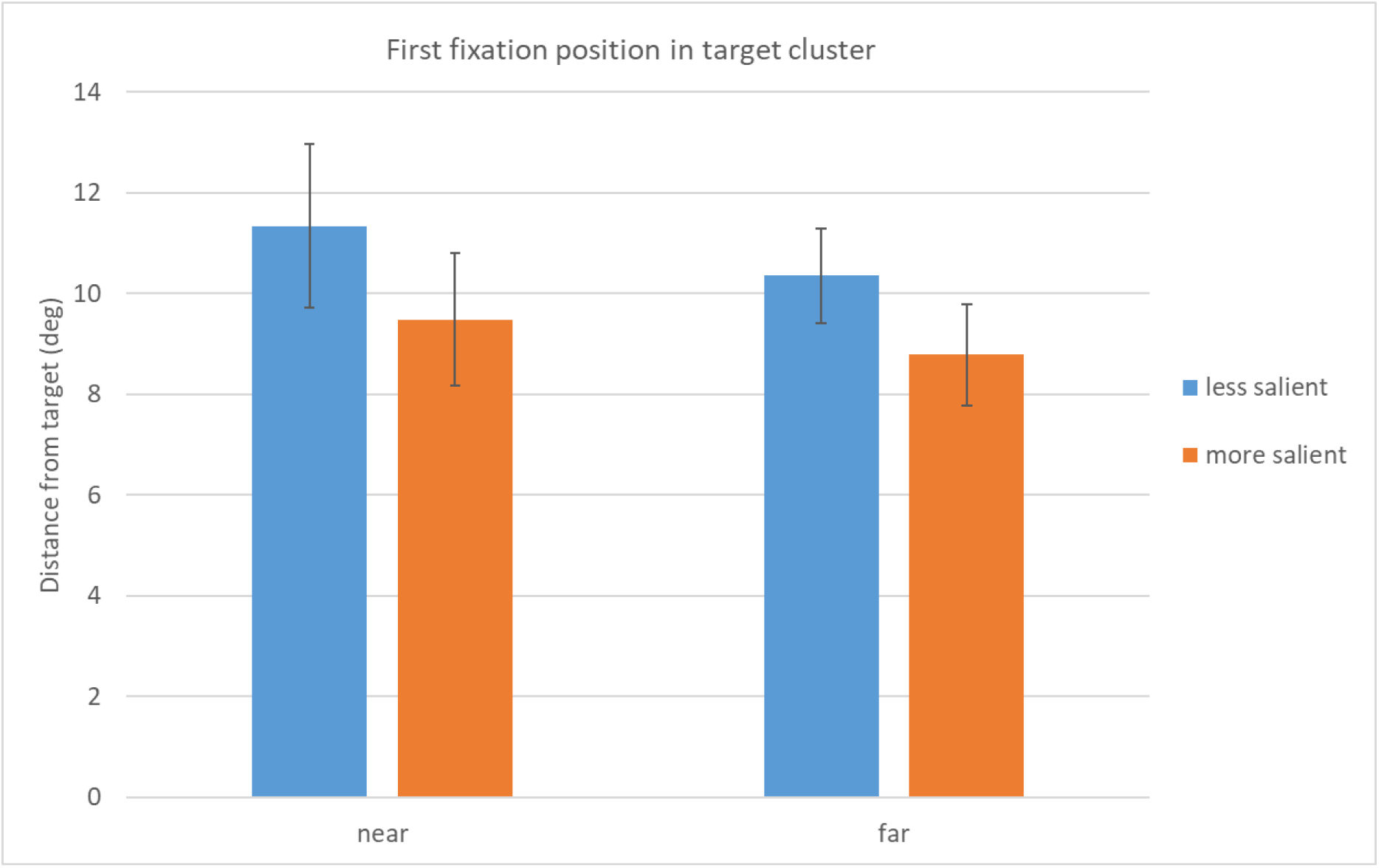
Distance between the first fixation in the target cluster and the target. N=23, error bars s.d. (note with a repeated measures comparison, error bars do not reflect significance). The less salient target is when distractors are oriented away from the target (horizontal or vertical) stimulus by +/- 20° and in the less salient condition only by +/-110°. Near and far are the cluster location relative to the fixation cross.

The polar plots show every single first fixation in a target cluster arranged so that polar plot angle is the angle away from the target relative to the centre of the cluster. We see that in general the fixations are more biased towards the centre of the cluster than elsewhere away from the target, but in the more salient condition at the far location it is clear that the bias is for fixations to be on target, whereas in the less salient condition at the far location it is towards the middle of the cluster. Again, if we compare the near to the far plots, we see a similar difference across both the more and less salient conditions. The first fixations in the target clusters are less spread away from the target in the far clusters and more centred around the direction of the centre of the target cluster

## Discussion

We designed two experiments where the task required participants to choose between and inspect up to four regions (clusters), in order to detect a target. The locations were arranged such that two were visible to participants when the trial began, whereas two required the participants to move their head to bring them into view. We manipulated the salience of the targets to investigate the way in which the location and salience of peripheral targets influences the planning of search strategy and associated landing positions of the saccades. In particular, we tested whether there was a difference in target detection in locations that became visible in the periphery by appearing at the start of trial versus those locations that only became visible in the periphery as the participants moved their eyes and heads to inspect the scene. Between the two experiments we varied the difficulty of the task. For the first experiment, once the target had been detected it was unambiguous. In the second experiment often some additional decision making was needed to select the target. This paradigm differs from 2D screen based visual search in that what is present in the observer’s FoV is changed by their head movement and eye movements are used to select locations within the new FoV.

In experiment 1 we found some interesting differences between the effect of salience on the time it took to reach the target cluster (i.e. change FoV so that the cluster was towards the centre and easily fixated), versus the effect of salience on the ability to target the landing position of a saccade (the accuracy of the first gaze position within the FoV centred on a new cluster). In the first experiment using letters as stimuli, we found that the more salient targets were detected more quickly when they appeared in the locations visible at the start of the trial (‘near’). This suggested that they were more often identifiable from the periphery and thus the first head and eye movements could be made in the correct direction. This was confirmed by 90% vs 58% of first saccades being in the correct direction for the more salient stimuli. However there appeared to be no advantage for salience at the far locations that came into view once the near locations were being inspected, suggesting that the far target had no influence in making the next movement in the correct direction or ‘abandoning’ the near cluster sooner to move to the far target. Somewhat surprisingly however, in both near and far locations the first saccade to land in the cluster containing the target landed closer to the more salient than the less salient target. Looking at Figure 5 we can see that this decrease appears to be due to more fixations being biased towards the target location on average. A further notable effect was that the first fixations in the cluster containing the target were closer to the target for the far than the near locations for both types of targets. Salience did not change this effect of location, and based on Fig 5 it can be seen that even in the salient conditions many fixations land on the centre of the cluster (i.e. there are fixations even in the salient case not aimed at the target), overall still resulting in an average decrease in distance between target and fixation, and are more tightly arranged around the centre of the cluster at the far locations.

A similar dissociation was found in the second experiment using Gabor patches. In this experiment there was no effect of the salience of the target on the time it took to reach a near target cluster (i.e. to change FoV so that the relevant cluster was towards the centre as it is first fixated), suggesting there was not enough information available in the periphery to make the first head and eye movement in the correct direction. This was confirmed by only 50% vs 48% of the first eye movements being in the correct direction for the more salient target in the near location. As before, there was also no advantage for salience at the far locations that came into view once the near locations were being inspected, suggesting that the far target had no influence in making the next movement in the correct direction or ‘abandoning’ the near cluster sooner to move to the far target. Yet even in this experiment, even though based on the time it took for the eyes to first reach the target cluster there was no effect of salience, in both locations the first saccade to land in the cluster containing the target landed closer to the more salient than the less salient target. Looking at Figure 8., again we can see that this decrease appears to be due to more fixations being biased towards the target location on average. However, again, reflecting the main effect of cluster location in Fig. 8 it can be seen that even in the salient conditions many fixations land on the centre of the cluster (i.e. there are fixations even in the salient case not aimed at the target). Overall this still results in an average decrease in distance between target and fixation, and fixations are more tightly arranged around the centre of the cluster at the far locations.

**Figure 8.**
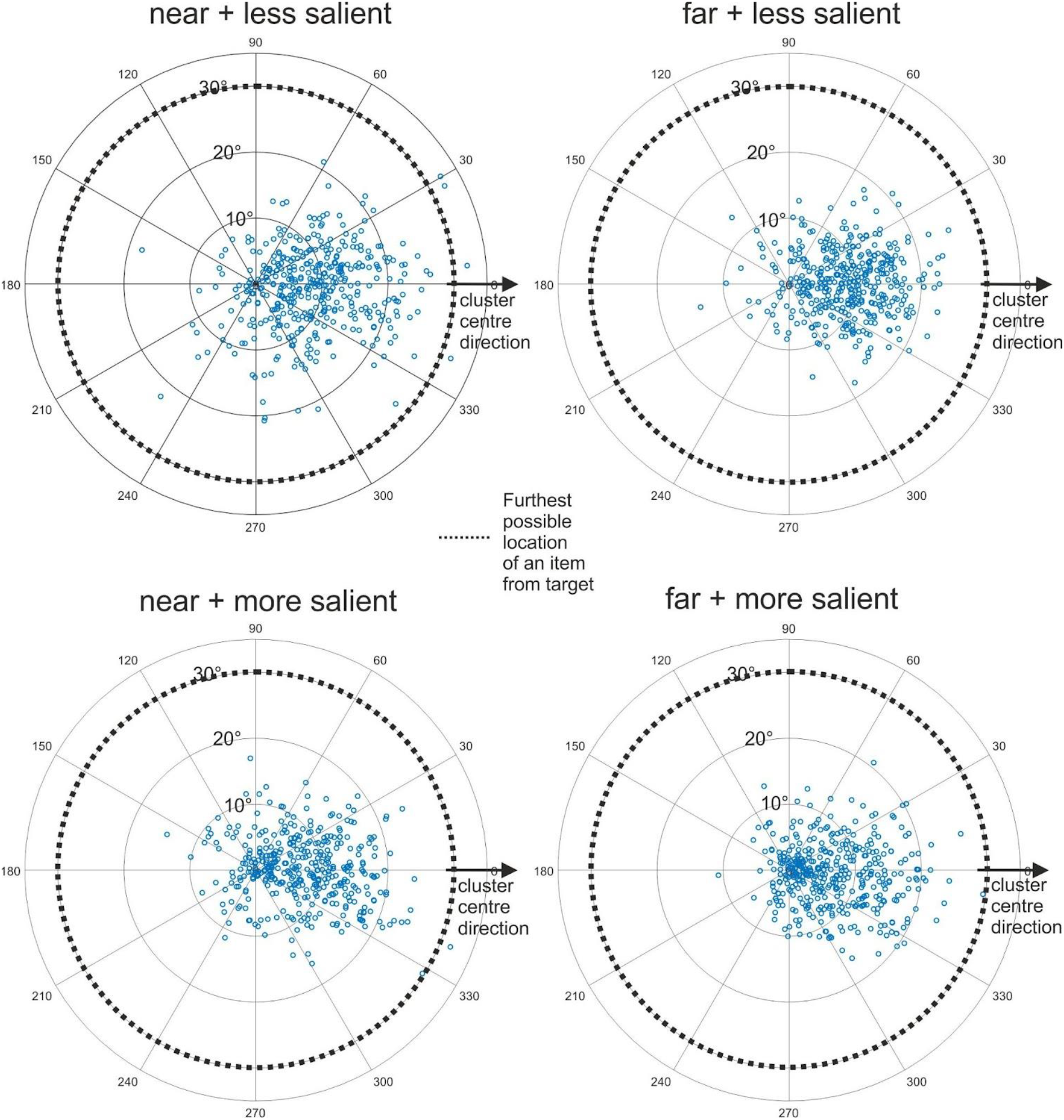
Polar plots of the first fixations in a cluster. The polar plots show every single first fixation in a cluster arranged so that the polar plot angle is the angle away from the target relative to the centre of the cluster.

This pattern of results suggests there are differences in visual search when a target appears in the field of view involuntarily versus based on the participants’ own actions. First, the effect of salience is not equivalent between the two target appearance types. The reaction time advantage for the more salient targets is completely absent when the target appears in the field of view through participant motion. This suggests that some information about target location can be available to eye movement planning that is not able to influence the global search strategy to speed up detection of more salient targets in the far locations. Second, there is a difference in fixation strategy according to how the target enters the FoV. In both experiments the far locations have more accurate fixations in the more salient conditions and more first fixations in the cluster that land towards the centre regardless of the salience of the target.

The peripheral location of the clusters when they enter the FoV, either by self-motion or involuntarily, raises the possibility of crowding. This is the effect whereby target stimuli presented in the periphery can be spatially masked by distractor stimuli placed in their proximity. Masking increases the further into the periphery the stimuli are presented and is greater than would be expected based on the peripheral stimulus magnification required for recognition of individual stimuli (Greenwood et al., 2009). It is also greater the more similar the surrounding objects are to the target (Wilkinson et al., 1997). Target stimuli presented at 35DVA into the periphery may suffer from crowding even if they have the capacity to pop out when presented closer to fixation. For instance, it is highly likely crowding occurred to the Gabor pattern stimuli. Some evidence has been presented to suggest that the act of making a saccade toward a peripheral stimulus can abolish crowding (Harrison et al., 2013). It is possible that this release from crowding combined with a head movement is what allows for the more accurate targeting of the salient stimulus. It has been suggested, however, that position information is lost in crowding (Greenwood et al., 2009). In our stimuli we see that although early identity information is not available always to guide the head movements toward clusters, for the more salient targets positional information is not always completely lost.

It seems there is some information available from the periphery about the location of the more salient targets that is not available for the less salient, allowing the eyes land more often near the more salient target, even in the difficult Gabor based task. This information, however, does not seem to influence in all cases the speed with which the eyes move to the target location. This implies that the location of the salient target within the cluster is represented, but the fact that the target is within the region of the cluster does not become available to the participant in a way which allows them to prioritise centring their FoV on the target cluster. This suggests a dissociation in the types of information available to different aspects of gaze location planning. Broad targeting of head movements that change the FoV may be driven by a pre-planned strategy that fails to register the more precise position information available to move the eyes to a location within the FoV. This reveals different levels of access to targeting information for different components of the search task in 360 degree space.

This pattern of results also suggests that first the head movement is made to shift the FoV and then the eye movement follows. Zangemeister & Stark (1982) suggested that under predictable and speeded conditions such as this visual search task it is more common for the head movement to occur first. Once the head is in motion in our experiments, the sequence of FOV shifts is not influenced by the salience of the target. Once the FOV does shift to the cluster containing the target, the landing location of the eye movement is influenced by the salience of the target. The head movement thus appears to be reliant on a pre-planned strategy. Spatial information is available to the eye movement and this supports greater precision of the eyes within a given FoV, but this information does not translate to greater accuracy of the head movement that adjusts the FOV.

Head and eye movements are commonly characterized as being synchronously planned based on the same visual information (André-Deshays et al., 1988; Barnes, 1979; Bizzi et al., 1971; Moschner & Zangemeister, 1993), however there are some examples to the contrary. Corneil & Munoz (1999) found occasional early head shifts toward an incorrect distractor that was in the opposite direction of the eye movement shift towards the correct target. Similar to our results this suggests correct visually acquired information is overridden by a rapid pre-planned movement. Ron et al. (1993) also found that when gaze shifts involved some element of pre-planning, eye movements could follow head movement and even become decoupled, suggesting the prioritisation of different locations by different types of orienting movement. Similarly, Morasso et al., (1973) suggest that saccades made during a head movement may be different to what is observed for saccades without; the former are not entirely predetermined by initial oculomotor commands. Both the experiments presented here also demonstrate different patterns for how the eyes land in the clusters depending on how they appear in view. The fixations that land in the near clusters are seemingly more randomly scattered than those in the far clusters, where a higher proportion of first fixations land towards the centre of the cluster. Thus, how an object comes into view does affect how we move our eyes towards it. One difference could be that gaze shifts away from fixation cross are based on the perceived location of their constituent items, whereas for the far clusters this could be based more on the prior representation of the average location of the clusters. However, the effect of salience remains the same, more salient targets are fixated more accurately, suggesting that it is the broad targeting of the head movement that changes between near and far target locations, rather the positional information available about the more salient stimulus.

Tatler & Land (2011) have suggested there is a less precise spatial memory based system for changing the field of view using head (and torso) movement and once this FoV has changed a more precise saccade can be made to the object of interest. What is interesting and somewhat surprising in our data is that the very first gaze location within a cluster seems to have access to the more precise position information of the salient target. This implies the more precise saccade is planned in concert with the FOV change, even though it does not influence when that specific FOV change will be executed. Further research in 360 degree and fully immersive virtual reality has the potential to tease apart the timing of planning and targeting of both types of head and eye movements.

In conclusion, our experiments show that there is indeed a difference between a target that comes in to view at onset versus one that comes into view due to a head movement and the pattern of differences suggests that although we may have spatial information in some cases for how to move our eyes within a potential new FoV, this information is not always consciously available to us when making the decision of which way to move our heads in order to change the FoV, thus revealing a somewhat surprising dissociation between these two levels of decision making that feed into our stream of everyday conscious experience.

## References

André-Deshays, C., Berthoz, A., & Revel, M. (1988). Eye-head coupling in humans: I. Simultaneous recording of isolated motor units in dorsal neck muscles and horizontal eye movements. Experimental Brain Research, 69(2). https://doi.org/10.1007/BF00247585

Barnes, G. R. (1979). Vestibulo-ocular function during co-ordinated head and eye movements to acquire visual targets. The Journal of Physiology, 287(1), 127–147. https://doi.org/10.1113/jphysiol.1979.sp012650

Bizzi, E., Kalil, R. E., & Tagliasco, V. (1971). Eye-Head Coordination in Monkeys: Evidence for Centrally Patterned Organization. Science. https://doi.org/10.1126/science.173.3995.452

Brysbaert, M. (2019). How many participants do we have to include in properly powered experiments? A tutorial of power analysis with reference tables. Journal of Cognition, 2(1), 16. https://doi.org/10.5334/joc.72

Corneil, B. D. (2011, August 18). Eye-head gaze shifts. The Oxford Handbook of Eye Movements. https://doi.org/10.1093/oxfordhb/9780199539789.013.0016

Corneil, B. D., & Munoz, D. P. (1999). Human Eye-Head Gaze Shifts in a Distractor Task. II. Reduced Threshold for Initiation of Early Head Movements. Journal of Neurophysiology, 82(3), 1406–1421. https://doi.org/10.1152/jn.1999.82.3.1406

Crundall, D. (2005). The integration of top-down and bottom-up factors in visual search during driving. In Cognitive processes in eye guidance (pp. 283–302).

Doshi, A., & Trivedi, M. M. (2012). Head and eye gaze dynamics during visual attention shifts in complex environments. Journal of Vision, 12(2), 9–9. https://doi.org/10.1167/12.2.9

Fang, Y., Nakashima, R., Matsumiya, K., Kuriki, I., & Shioiri, S. (2015). Eye-Head Coordination for Visual Cognitive Processing. PLOS ONE, 10(3), e0121035. https://doi.org/10.1371/journal.pone.0121035

Greenwood, J. A., Bex, P. J., & Dakin, S. C. (2009). Positional averaging explains crowding with letter-like stimuli. Proceedings of the National Academy of Sciences, 106(31), 13130–13135. https://doi.org/10.1073/pnas.0901352106

Harrison, W. J., Mattingley, J. B., & Remington, R. W. (2013). Eye Movement Targets Are Released from Visual Crowding. Journal of Neuroscience, 33(7), 2927–2933. https://doi.org/10.1523/JNEUROSCI.4172-12.2013

Hulleman, J., & Olivers, C. N. L. (2017). The impending demise of the item in visual search. Behavioral and Brain Sciences, 40. https://doi.org/10.1017/S0140525X15002794

Morasso, P., Bizzi, E., & Dichgans, J. (1973). Adjustment of saccade characteristics during head movements. Experimental Brain Research, 16(5), 492–500. https://doi.org/10.1007/BF00234475

Moschner, C., & Zangemeister, W. H. (1993). Preview control of gaze saccades: Efficacy of prediction modulates eye-head interaction during human gaze saccades. Neurological Research, 15(6), 417–432. https://doi.org/10.1080/01616412.1993.11740176

Olk, B., Dinu, A., Zielinski, D. J., & Kopper, R. (2018). Measuring visual search and distraction in immersive virtual reality. Royal Society Open Science, 5(5), 172331. https://doi.org/10.1098/rsos.172331

Ron, S., Berthoz, A., & Gur, S. (1993). Saccade-vestibulo-ocular reflex co-operation and eye-head uncoupling during orientation to flashed target. The Journal of Physiology, 464(1), 595–611. https://doi.org/10.1113/jphysiol.1993.sp019653

Shioiri, S., Kobayashi, M., Matsumiya, K., & Kuriki, I. (2018). Spatial representations of the viewer’s surroundings. Scientific Reports, 8(1), 7171. https://doi.org/10.1038/s41598-018-25433-5

Stern, J. A., Brown, T. B., Wang, L., & Russo, M. B. (2005). Eye and head movements in the acquisition of visual information. Psychologia, 48(2), 133–145. https://doi.org/10.2117/psysoc.2005.133

Tatler, B. W., & Land, M. F. (2011). Vision and the representation of the surroundings in spatial memory. Philosophical Transactions of the Royal Society B: Biological Sciences, 366(1564), 596–610. https://doi.org/10.1098/rstb.2010.0188

Vater, C., Williams, A. M., & Hossner, E.-J. (2020). What do we see out of the corner of our eye? The role of visual pivots and gaze anchors in sport. International Review of Sport and Exercise Psychology, 13(1), 81–103. https://doi.org/10.1080/1750984X.2019.1582082

Wilkinson, F., Wilson, H. R., & Ellemberg, D. (1997). Lateral interactions in peripherally viewed texture arrays. JOSA A, 14(9), 2057–2068. https://doi.org/10.1364/JOSAA.14.002057

Wolfe, J. M. (2020). Visual Search: How Do We Find What We Are Looking For? Annual Review of Vision Science, 6(1), 539–562. https://doi.org/10.1146/annurev-vision-091718-015048

Zangemeister, W. H., & Stark, L. (1982). Types of gaze movement: Variable interactions of eye and head movements. Experimental Neurology, 77(3), 563–577. https://doi.org/10.1016/0014-4886(82)90228-X

